# Oriented Cell Dataset: efficient imagery analyses using angular representation

**DOI:** 10.1101/2024.04.05.588327

**Authors:** LN Kirsten, AL Angonezi, FD Oliveira, JL Faccioni, CB Cassel, DC Santos de Sousa, S Vedovatto, CR Jung, G Lenz

**Affiliations:** Institute of Informatics, Federal University of Rio Grande do Sul (Brazil); Institute of Biosciences, Federal University of Rio Grande do Sul (Brazil)

**Keywords:** Cell detection, oriented bounding boxes, cell morphology, deep learning

## Abstract

In this work, we propose a new public dataset for cell detection in bright-field microscopy images annotated with Oriented Bounding Boxes (OBBs), named Oriented Cell Dataset (OCD). We show that OBBs provide a more accurate shape representation compared to standard Horizontal Bounding Boxes (HBBs), with slight overhead of one extra click in the annotation process. Our dataset also contains a subset of images with five independent expert annotations, which allows inter-annotation analysis to determine if the results produced by algorithms are within the expected variability of human experts. We investigated how to automate cell biology microscopy images by training seven popular OBB detectors in the proposed dataset, and focused our analyses on two main problems in cancer biology: cell confluence and polarity determination, the latter not possible through HBB representation. All models achieved statistically similar results to the biological applications compared to human annotation, enabling the automation of cell biology and cancer cell biology microscopy image analysis. Our code and dataset are available at https://github.com/LucasKirsten/Deep-Cell-Tracking-EBB.

## I. Introduction

Detection and tracking of living cells in microscopy images is a crucial task required in many medical and research applications, such as cell growth, migration, invasion, morphological changes, and alterations in the localization of molecules within cells [1]–[4]. Bright-field microscopy has several advantages, such as not requiring any fluorescent tagging of the cell, reduced photo-toxicity, and much more affordable microscopes [5]. The drawback of bright-field images is the difficulty of automating analysis due to the lower contrast to the background compared to fluorescence microscopy, and images might contain artifacts similar to the cells. Multiple approaches have been proposed to detect cells in bright-field microscopy images [6], but the diversity of microscopes, cell lines, and level of magnification hinders the possibility of applying the same pipeline to different experimental setups.

Besides identifying cells, measuring their size and shape is relevant for identifying phenotypes such as senescence [7] and epithelial to mesenchymal transition (EMT) [8]. Horizontal Bounding Boxes (HBBs) are the *de facto* choice for generic object detection [9] but they are not suited to obtain the actual shape and size of the cells (see Fig. 1a). Obtaining the full cell masks through segmentation approaches is an alternative, but the annotation process is tedious and ill-defined for overlapping cells [10] (see Fig. 1a) or low-contrast images (see Figs. 2c and 2d). We advocate that Oriented Bounding Boxes (OBBs) are adequate representations for cellular imagery applications, with a good compromise between object representation and annotation cost. Compared to HBBs, they require an additional angular parameter, allowing a much better representation of the cell shape (see Fig. 1b). In particular, HBBs might contain large portions of background and are not able to capture the aspect ratio of elongated orientated cells, which compromise their use in applications such as cell confluence estimation (relative area occupied by the cells) and polarity (ratio between the major and minor axis of the cell shape), as illustrated in Fig. 1. As an additional advantage, the orientation/shape of detected cells can be explored in tracking-by-detection approaches to provide a better spatio-temporal association [11].

**Fig. 1:**
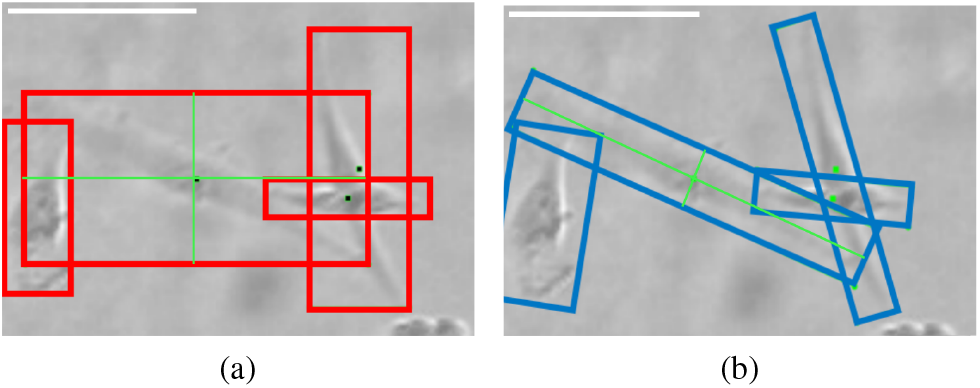
Comparison of (a) HBB and (b) OBB annotations for glioblastoma cells. Green lines show bounding box axes, which are used to calculate the cell area and polarity. For the middle cell, the area is *∼*2.6 times larger whereas the polarity is *∼*3 times smaller when HBB and OBB of the same cell are compared. Scale bar (top-left white) equals 30 *μ*m.

**Fig. 2:**
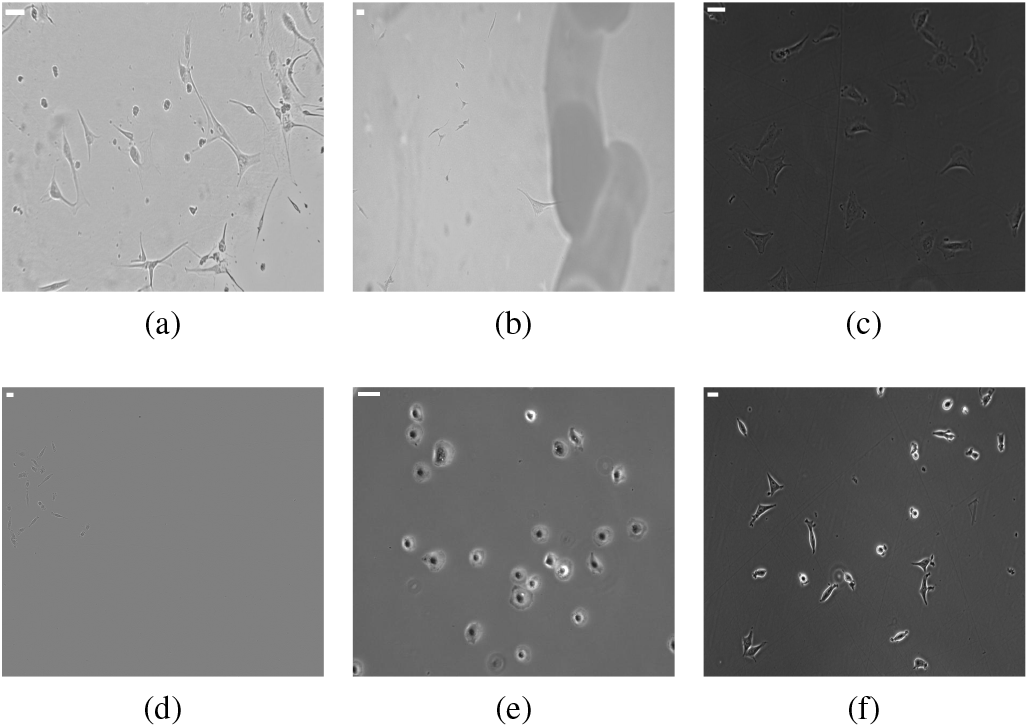
Example of different images from the OCD. (a) and (b) are A172 cells captured in CytoSMART microscope; (c) and (f) are MRC5 and MCF7 cell lines, respectively, captured in Zeiss Axiovert 200 microscope; (d) are U251 cells captured in IncuCyte; and (e) are SCC25 cells captured in a Zeiss AxioCam microscope. Scale bar (top-left white) equals 30 *μ*m.

In this work, we advocate that OBBs are suitable for cell representation, and propose a new public “Oriented Cell Dataset” that provides annotated bright-field cell images as oriented bounding boxes. We also investigate using OBBs for cell detection in bright-field microscopy images, focusing on two important biological applications: cell confluence, and polarity estimation. Our main contributions are: i) we propose a publicly available “Oriented Cell Dataset” that provides annotated bright-field cell images as oriented bounding boxes; ii) using the popular Cell Tracking Challenge (CTC) [12], we show that oriented representations such as oriented bounding boxes (OBBs) and oriented ellipses (OEs) present considerable more overlap with the segmentation masks than traditional horizontal bounding boxes (HBBs); iii) we perform an inter-annotator assessment (IAA) evaluation of OBB human annotations to estimate the human variability, and propose a suitable IoU threshold for evaluating automatic OBB cell detectors; iv) we show that oriented object detectors can be employed to automate biological tasks of cell detection, confluence and polarity estimation.

## II. Related work

There are several approaches for cell counting, detection, and segmentation in bright-field microscopic images, the best results being achieved by deep learning methods [13]. These methods vary considerably regarding the network architecture and the *degree of supervision* required to label training data: detecting just the cell center requires one pixel-per-cell as supervision; detecting the cell boundaries as an HBB requires two points (top-left and bottom-right), while OBBs requires an additional parameter related to the orientation; finally, segmentation requires the identification of all pixels belonging to a cell, which is very time-consuming [6].

As noted in [6], a popular approach for cell detection in microscopy images is to use specialized versions of algorithms for general-purpose object detection/segmentation, such as Faster-RCNN [14]. General-purpose object detection with HBBs is already a well-consolidated problem in computer vision with several improvements in recent years, as noted in the recent survey paper [9]. However, the representation of cells as HBBs presents limitations in denser scenarios, particularly when oriented and elongated cells are present, as illustrated in Fig. 1a. Meanwhile, the literature regarding object detection with OBBs is recent and scarce, and focuses mostly on niche applications such as aerial/satellite imagery [15], [16]. Although HBB and OBB detectors share the same main concepts in terms of network architecture (e.g., both can be achieved using two- or one-stage methods), moving from HBBs to OBBs adds some challenges, such as the ambiguous parametrization of OBBs, the difficulty in regressing angular information, and the adaptation of anchor-based methods [16].

In two-stage methods, the first stage creates OBB proposals, and the second predicts the class-related confidence for each proposal and refines its shape. For example, Xia et al. [15] adapted the popular Faster-RCNN method [14] (renamed as RetinaNet) to generate and evaluate OBB proposals in the context of object detection in aerial/satellite images. To avoid developing specific pooling strategies for OBBs, R2CNN [17] explores three horizontal RoI-pooling layers on the enclosing box of the OBB, and develops a regression strategy for generating OBBs as final representations.

One-stage methods aim to simultaneously regress an OBB related to an object and predict its class label and are more popular than two-stage methods nowadays. Yang et al [18] proposed R3Det, which focuses on refining OBBs through pixel-wise feature interpolation. Qian et al [19] proposed the RSDet model, which uses the RetinaNet [20] architecture with a new loss combined with an eight-parameter regression method (instead of the usual five-parameter) to solve the problem of inconsistent parameter regression in OBBs. To solve the discontinuous boundaries issue originated by the angular periodicity or corner ordering, Yang et al. proposed the Circular Smooth Label model (CSL) [21] and the Densely Coded Labels model (DCL) [22], which transform the angular prediction task from a regression to a classification problem. Although these methods have been broadly used for detecting oriented objects in aerial/satellite images [15], [16], we are unaware of existing datasets that provide OBB annotations for bright-field microscopic images. As mentioned before, using OBBs might severely reduce the annotation burden when compared to segmentation approaches, and still provide enough geometrical information to enable cell confluence and polarity estimation, which are broadly used on biological applications [1]–[3], [8]. In this work, we evaluate the use of such models for detecting cells in bright-field microscopic images by comparing their detections with human annotators, and with standard literature metrics used for general-purpose object detection applications using our proposed Oriented Cell Dataset.

Current open-source datasets for cell detection usually provide annotations as single-marker points [23] or as HBBs [24], which are not suitable for cell polarity or confluence estimation. Meanwhile, the most popular benchmark for cell segmentation and cell tracking, the Cell Tracking Challenge (CTC) [12], uses computer-origin reference annotations to annotate most of the cell masks (only 5.37% of the cells are annotated by humans). These examples highlight the difficulties in providing high-quality biological datasets with human annotations. In this work, we propose a public “Oriented Cell Dataset” that provides fully human annotations of bright-field cell images as OBBs. We also provide a smaller subset of images annotated by five different human experts and perform a quantitative analysis of inter-human annotation agreement. As noted in [25], even small discrepancies in bounding box annotations can lead to strong IoU degradation. In this work, we show that this behavior is amplified in cell microscopy imagery due to the inherent difficulty of the problem. Finally, we demonstrate that this dataset can be used with oriented detectors to automate biological tasks of cell detection, polarity, and confluence estimation.

## III. Oriented Cell Dataset (OCD)

We created a new public cell dataset^1^, called *Oriented Cell Dataset* (OCD) [26]^2^, containing OBB annotations for bright-field microscopy images. A total of 160 images were acquired using different microscopes and cell lines, which resulted in visually distinct images as shown in Fig. 2. These images were split into three sets:

- **Train**: 120 images (75% of total) used for training the object detectors;
- **Test**: 30 images (18.75% of total) used for evaluating the detectors performance, and
- **Annotator’s Comparison**: 10 images (6.25% of total) used for the inter-annotator assessment and biological applications validation (see Sec. IV-B).

Each image of the *Train* and *Test* splits was annotated by a single annotator, and they contain an equal distribution (stratified) of their differences regarding the used microscope, cell lines, and cultivation method. Meanwhile, each image of the *Annotator’s Comparison* split was annotated by five different human experts. This last split also contains images from two microscopes (IncuCyte and Zeiss AxioCam) and one cell line (human squamous cell carcinoma) that were not present in the previous splits to evaluate the generalization capabilities of the trained models. A full description of the dataset acquisition and splits is described in Tab. I.

**TABLE I:**
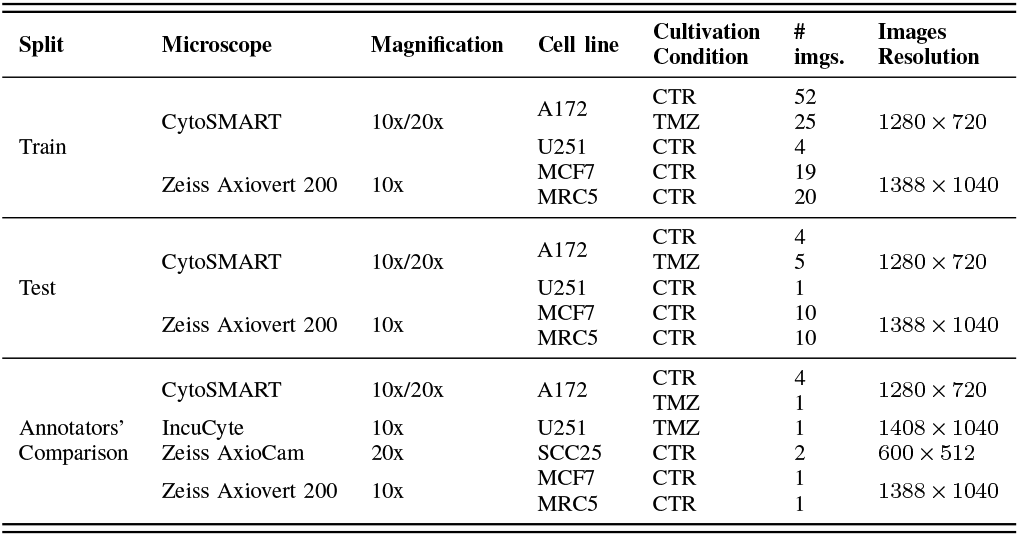
OCD description. Cell lines A172 and U251: human glioblastoma; MCF7: human breast cancer; MRC5: human lung fibroblast; SCC25: human squamous cell carcinoma. Cultivation condition CTR: cells belonging to the control group - without the addition of chemotherapy; TMZ: cells treated with 50 *μ*M temozolomide in some cultivation step - 3h treatment in most experiments.

All images were manually annotated using the roLabelimg tool^3^, which allowed the researchers to delimit the OBBs (composed of *x*-center, *y*-center, height, width, and angle) for each cell. Furthermore, the cells were classified either as “elongated” or “round”, i.e. mitotic, cells, as illustrated in Fig. 3. In total, over 6,700 OBBs were annotated, from which 89.4% were classified as “elongated” and 10.6% as “round” cells. The confluence (i.e., the area occupied by cells in a given image *I*) was computed as:

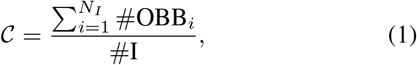

where #OBB_*i*_ is the OBB area of cell *i, N*_*I*_ is the number of cells in image *I*, and #I is the image area. We obtained a mean confluence among the images of *C ≈*10%, ranging from nearly 0% to 40%.

**Fig. 3:**
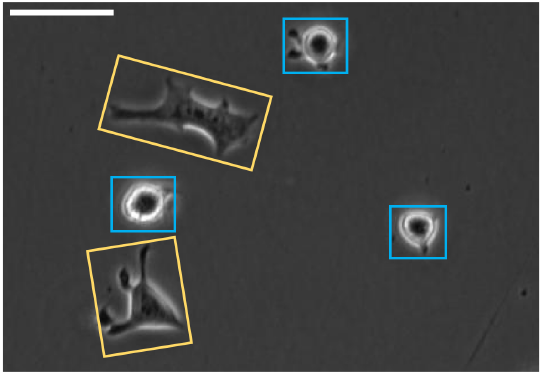
Illustration of “elongated” (yellow) and “round” (blue) cells in the OCD. Scale bar (top-left white) equals 30 *μ*m.

## IV. Experiments and Applications

In our experimental setup, we first evaluate if OBBs are suitable approximations for the cell shape. Then, we perform an inter-annotator assessment (IAA) experiment to evaluate the degree of agreement among the human annotators and discuss suitable acceptance thresholds. Using the *Train* split of our dataset, we train several object detectors to asses if their variability is similar to what is expected among human annotators. Finally, we illustrate the usefulness of OBB cell detection in two biological applications: cell confluence and polarity estimation.

### A. Are OBBs adequate representations?

To evaluate if OBBs are good approximations for the cell shapes, we evaluated seven public datasets from the Cell Tracking Challenge (CTC) [12]^4^. These datasets contain annotated segmentation masks, provided as *silver* and *gold* standards. The silver standard annotations refer to computer-origin reference annotations, while the gold standard refers to human-origin ones. Since only 5.37% cells are annotated in the gold standard, most works use only the silver standard annotations. This emphasizes the difficulty of manually annotating segmentation masks on microscopic images.

We considered all provided segmentation masks (silver and gold standards) as the reference (ground-truth) cell shape representation and fitted both HBB and OBB representations as approximations of the mask based on provided OpenCV^5^ implementations. We also considered the natural extensions of these approximations for fitting elliptical-like objects: converting an HBB to an Axis-Aligned Ellipse (AAE) with major and minor axis as the major and minor HBB lengths; and the OBB to an Oriented Ellipse (OE) with major and minor axis as the major and minor OBB lengths, respectively. Both center points and angle of rotation (in the OBB case) are kept the same in the AAE and OE representations. The four considered approximations were evaluated with the Intersection over Union (IoU), computed as:

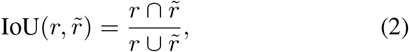

where *r* is the reference shape representation (i.e., the segmentation mask), 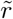 is the approximate representation (HBB, OBB, AAE or OE). Furthermore, we computed the average of the minimal number of points required to annotate the cells of each dataset using the segmentation masks to evaluate the annotation efforts.

### B. Inter-annotator assessment

In generic-purpose object detection, the presence, category, and HBB/OBB for each instance are typically well-defined. On the other hand, annotating biomedical data is far more challenging, and even experts might disagree on some data (see Fig. 8). As such, the ground-truth annotations are expected to comprise a “human variability” factor [25], which can also impact the objective metrics used to compare object detection approaches [27]. In this section, we propose to evaluate the level of agreement among human annotators regarding cell detection and classification (a.k.a. Inter-Annotator Assessment, or IAA).

We used a smaller set of ten images that were independently annotated by five researchers (having multiple annotators for the full dataset would be unfeasible), composing the *Annotators’ comparison* data split (recall Sec. III). Inspired by [28], we explore the Krippendorff’s Alpha (*K −α*) metric for IAA. As in [28], we tackle the problem from the semantic segmentation perspective, where each image pixel presents a single label within a set of *X* values inherited from the OBB annotations. We propose two different categorizations in our analysis: i) a *class-aware* version, where each pixel is labeled as background, elongated, or round cell (*X* = 3); and ii) a *class-agnostic* version, where each pixel is labeled as either background or cell (*X* = 2). For more details regarding *K −α*, we refer the reader to [28].

The Krippendorff’s Alpha score provides an overall agreement metric among the annotators, but the score might be dominated by background pixels. Furthermore, it is not able to distinguish overlapping cells. On the other hand, the agreement of the OBBs annotated by two experts for the same cell is crucial to evaluate the precision, recall, AP, and mAP metrics, which are the standard metrics for generic-purpose object detection [29] and can be strongly affected by choice of the IoU threshold [27]. To evaluate the individual effect of cell shape annotation, we performed a second experiment with the *Annotators’ comparison* data split.

More precisely, let us consider *𝒜*_*i*_ and *𝒜*_*j*_ the set of annotated OBBs related to annotators *i* and *j*, respectively, and let *N*_*i*_ and *N*_*j*_ denote the corresponding number of annotated cells. For each pair of annotators *i ≠ j*, we consider *𝒜*_*i*_ as the GT and *𝒜*_*j*_ as the set of candidate detections. Since there is no confidence associated with the OBBs, we cannot compute the mAP metric [29]; instead, we use the F1-Score to evaluate the detection set, varying the IoU threshold (*T*) to validate a candidate detection. Since our goal is only to evaluate the geometric consistency of the OBBs, we performed the analysis in a class-agnostic manner only (i.e., both elongated and circular cells are considered in the same category).

Since typically *N*_*i*_ *≠ N*_*j*_, an analysis with the raw (unfiltered) annotations provides the joint effect of *T*, false positives (FPs), and false negatives (FNs). To isolate the individual effect of the threshold *T*, we have also performed an analysis with a filtered set of detections, obtained by applying the Hungarian Algorithm between every pair of annotation sets *𝒜*_*i*_ and 𝒜_*j*_ using 1 *−* IoU as the association cost. The optimal association patch matches min*{N*_*i*_, *N*_*j*_*}* pairs of OBBs, but it might match OBBs with very small (or no) intersections that probably relate to different cells. To avoid such mismatches, we eliminated pairs of matched OBBs with an IoU smaller than 0.1.

### C. OBB detectors for automatic cell detection

We trained seven popular OBB detectors using the *Train* split of OCD. To allow reproducibility, we leverage the implementations available at AlphaRotate^6^ rotation benchmark to train using OCD. The chosen OBB detectors (in chronological order) are: R2CNN [17], RetinaNet [20], R3Det [18], RSDet [19], CSL [21], DCL [22], and R3Det DCL [18], [22].

As in works that use aerial images [15], [16], our cell images are usually too large for training the detectors directly. Hence, we first cropped those images to 512 *×* 512 patches with a 100-pixels stride, resulting in a total of 5,590 patches of images to train our models. All models used the ResNet50 [30] backbone with pre-trained weights on the ImageNet dataset [31]. For training the models, weight decay and momentum are set to 10^*−*4^ and 0.9, respectively. We employed Momentum Optimizer over 1 GPU and one image per mini-batch. All models were trained for 100,000 steps. The learning rate started at 10^*−*3^ and was reduced tenfold at steps 60,000 and 80,000, respectively. Finally, in all experiments, we applied random rotation and flips to augment the training data.

In order to evaluate the trained models, we used the mean average precision (mAP) [29]. This metric relies on the IoU between annotated and predicted cells, which involves an acceptance threshold. In this work, we selected this acceptance threshold as 0.5 based on the IAA experiment, as described in Sec. V-B.

### D. Biological applications

Object detection is typically not a goal but an intermediary result from which relevant information can be extracted. The estimation of cell confluence in a given image is a crucial task in migration, wound-healing [2], [3] and cell-adherence [4] assays. The detection of remodeling in cell morphology also poses an important role in the determination of cancer invasiveness since cell polarity changes are one of the indicators of the EMT [8]. All these applications can benefit from OBB object detection, but the result might differ: while OBBs provide an exact value for the cell polarity, they produce an approximation for confluence. Furthermore, it is important to consider the IAA analysis when interpreting these evaluation metrics.

To determine whether the trained models produce results within the expected human variability, we performed statistical analyses on the *Annotator’s comparison* data split, regarding two especially relevant tasks related to cell and cancer biology: determining the cell confluence and polarity, as such:

- **Confluence**: refers to the relative area occupied by cells in a given image as defined in Eq. 1;
- **Cell polarity**: refers to the ratio between the major and minor OBB sides for a given cell (see Fig. 1).

The Analysis of Variance for Randomized Block Design (ANOVA-RBD) test was applied for cell confluence estimation. The Kruskal-Wallis test was applied to the distributions of cell polarities. Both tests aim for the same assessment: to identify whether at least one of the analyzed groups is statistically different from the others. The first is a parametric test, applied to normal distributions, and the latter is a non-parametric test, which can be applied to non-normally distributed data, such as cell polarity (more details will be provided in Sec. V-D.2). Since the ANOVA-RBD enables us to distinguish the intra-group (same annotator, different images) and inter-group (same image, different annotators) variability, we were able to determine whether the variation in cell count and confluence estimation was significant, based on the inter-group statistic. For the Kruskal-Wallis tests, regarding cell polarity estimation, we performed a test for each image distribution (i.e., annotators and models polarity histograms). An *α* = 0.05 was used for both tests to determine statistical significance.

We also assessed the models performance on biological applications using the *Test* data split. Cell confluence estimation generates a single quantity, and the Mean Relative Error (MRE) was used as a quality metric. Polarity estimation was performed for each cell, and each image was characterized by the frequency distribution (histograms) of all cells. For this task, we used the chi-square distance to compare the histograms. Based on the chosen evaluation metrics, we can characterize model performance related to the expected human variability (with ANOVA-RBD and Kruskal-Wallis tests) and to the ground-truth data.

## V. Results

### A. Comparison of cell representations

We present the results for representation comparison experiments described in Sec. IV-A with the public CTC [12] datasets in Tab. II. We can observe that for all datasets the OBB representation significantly improves the shape estimation error when compared to both HBB and AAE. Specifically, the OBB representation provides 20.7% better shape representation than HBBs on average, and 4.7% better than the AAEs. When using OEs, the improvement is even higher: 37.7% when compared to HBB, and 19.3% to AAE. This can be related to the fact that cells usually present a more “round-like” shape, similar to a rotated ellipse. Regarding the annotation efforts of segmentation masks, we can observe that, while the OBB (and OE) representations usually require 3 points (clicks from the user on the screen to annotate), the masks can scale this number from 27 up to 405 points (last column of Tab. II), which corresponds to 9 to 135 times more labor required from the human experts. Hence, we advocate that cells can be annotated using OBBs, but the analysis can be performed using OEs for a better overlap with the segmentation masks.

**TABLE II:**
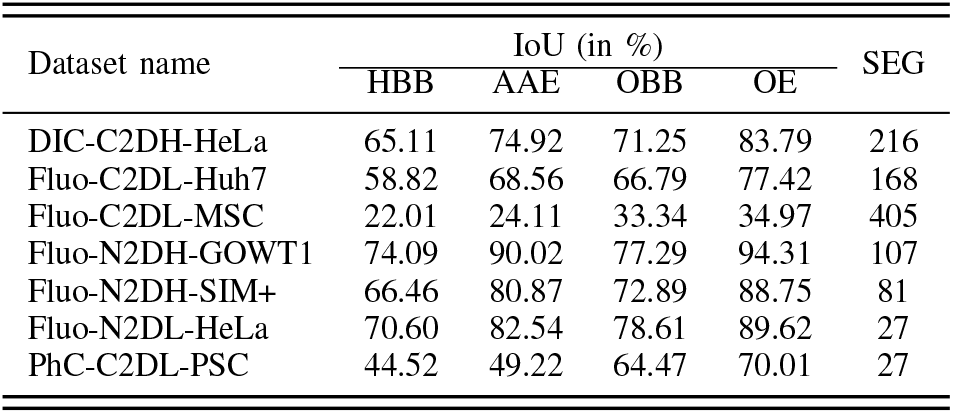
Comparison of different representations for approximating the segmentation masks on the CTC [12] datasets. The first four columns contain the IoU, and the last column (SEG) refers to the average minimal number of points required to define one segmentation mask on the dataset images.

### B. Inter annotator assessment

Following the experimental setup described in Sec. IV-B, we initially computed the *K −α* for the five human annotators. In the class-aware experiment, we obtained *K −α* = 0.760; in the class-agnostic experiment (where all cell annotations were considered as the same category) the *K −α* increased to 0.794. As noted in [32], a *K −α* value above 0.67 is considered good for IAA in the industry and academia, and above 0.8 is very good. Hence, we conclude that the agreement among the human annotators is between good and very good for both class-aware and class-agnostic experiments.

The second analysis described in Sec. IV-B focuses on the impact of the IoU threshold over the precision and recall metrics, consolidated through the F1-Score. We consider all pairwise combinations of annotators using both the raw (unfiltered) OBBs and the filtered version that considers only OBBs that were marked by both annotators.

Fig. 4 shows the plot of the mean F1-Score as a function of the IoU threshold (*T*) for the two scenarios (unfiltered and filtered), and the shaded regions indicate the variability when changing the pairs of compared annotators. As seen, the peak average F1-Score for the unfiltered experiments is around 0.8, which can be explained by FPs and FNs that are detected due to disagreements regarding what cell, debris, or background is. On the other hand, the results with the filtered annotations evaluate the individual effect of *T* for comparing the OBB of cells that were effectively marked by both experts but still might generate discrepancies in the shape and location. As we can see in Fig. 4, the results are very consistent for lower IoU thresholds but rapidly decay when more restrictive thresholds are used. This result clearly indicates that using the widely adopted COCO metrics [29], which considers a range of IoU values from 0.5 to 0.95, *is not realistic* for the cell detection problem using OBBs.

**Fig. 4:**
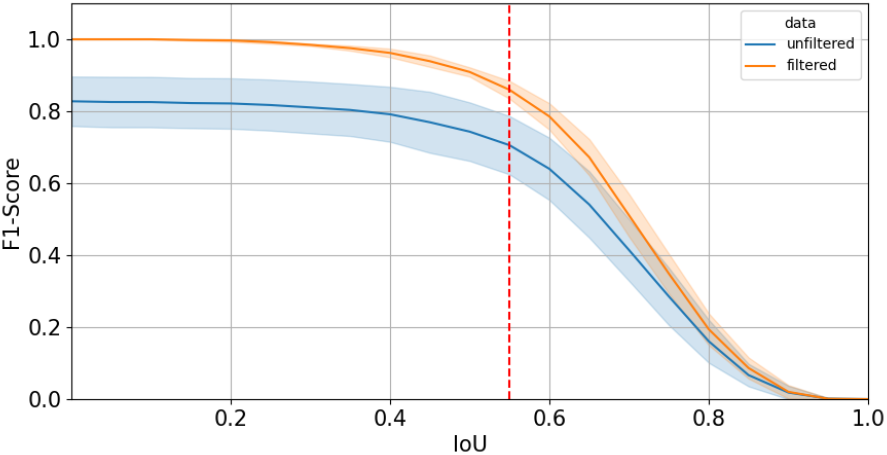
IAA evaluation based on F1-Score in different IoU thresholds. Dark lines refer to F1-Score mean values, shadow area indicates standard deviation, and the vertical dashed red line intersection with the curves refers to the curve’s knee-points.

To provide an IoU threshold that considers the inherent annotation variability, we computed the IoU value for which the relative cost of increment is no longer worth the corresponding performance benefit (i.e., the IoU value starts to be so restrictive that harms the F1-Score between pairs of human annotators) by finding the “knee-point” for the unfiltered and filtered data^7^ [33], which both arise around a threshold of *T* = 0.55. Since this value was obtained based on expert annotations, we suggest using it as *an upper bound* for oriented cell detectors. In fact, our findings are more aligned with the single threshold of 0.5 suggested in the Pascal VOC Challenge for object detection [34], which is used to evaluate the trained detectors next.

### C. Cell detection results

Based on the IAA analysis, we compare the seven trained OBB detectors using *T* = 0.5 for the main IoU threshold, as shown in the first column of Tab. III. We observe that R2CNN (a two-stage detector) achieves the highest mAP value using the suggested IoU threshold, and that R3Det (a one-stage detector) achieves the second highest. Nevertheless, we can observe that most detectors produced similar quality metrics, and that the least performing detector (CSL) resulted in a 14% lower mAP score than R2CNN.

**TABLE III:** Results in the *Test* data split. Cell detection was measured using the mAP_50_ [29], confluence error with the mean relative error (MRE), and polarities using the Chi-distance. Best values are in **bold**, and second-best <u>underlined</u>.

| Model | Cell detection ↑ | Confluence ↓ | Polarities ↓ |
| --- | --- | --- | --- |
| CSL | 63.20 | 0.104 | 2.66 |
| DCL | 67.58 | 0.099 | 2.61 |
| R2CNN | **73.49** | 0.083 | 2.27 |
| R3Det | 70.95 | 0.144 | 3.08 |
| R3Det DCL | 69.06 | 0.106 | 2.31 |
| RSDet | 68.97 | **0.072** | <u>2.21</u> |
| RetinaNet | 65.86 | <u>0.075</u> | **2.02** |

### D. Biological Applications of OBB Cell Detection

#### 1) Confluence estimation

By employing OBBs instead of HBBs, the estimation of the area occupied by a cell is substantially improved (recall Tab. II), enabling confluence estimation. To assess whether the trained models yielded results within human variability, an ANOVA-RBD test was applied. All models achieved area estimation values within human variability (*α* = 0.05), demonstrating the capability of using the presented models in experiments that require confluence estimation.

Fig. 5 shows the per-image confluence estimation produced by each method of the *Test* data split, along with the annotated ground-truth value. We can observe that lower-density images (4–5, 7–16, and 27–30) tend to produce the smallest errors. The confluence errors tend to grow as the cell density increases, which might actually be expected: as noted in [9], dense and occluded object detection is still challenging scenarios for generic-purpose object detection. The MRE considering all images in the set for all tested methods is shown in the second column of Tab. III. Most models presented MREs of *∼*10%, with RSDet, RetinaNet, and R2CNN, respectively, presenting the best results. As for the cell count estimation problem, we note that the best model for confluence estimation is not the same considering object detection metrics. In fact, R2CNN had an error *∼*15% higher than RSDet.

**Fig. 5:**
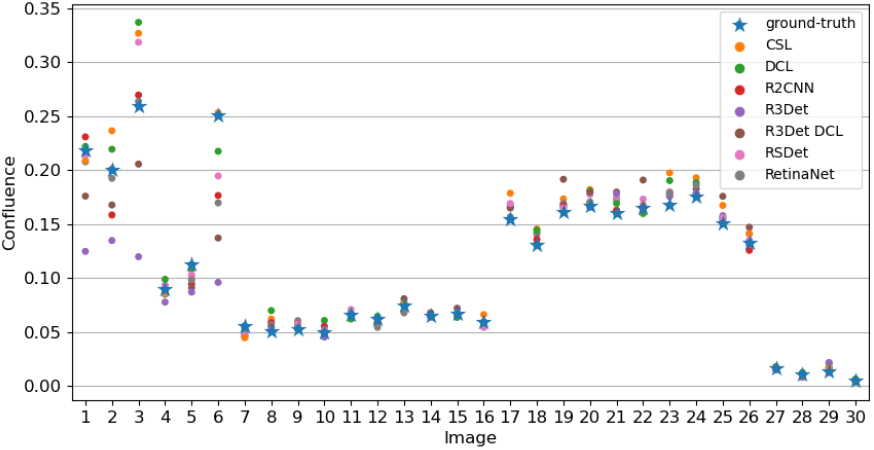
Cell confluence comparison on the *Test* dataset.

**Fig. 6:**
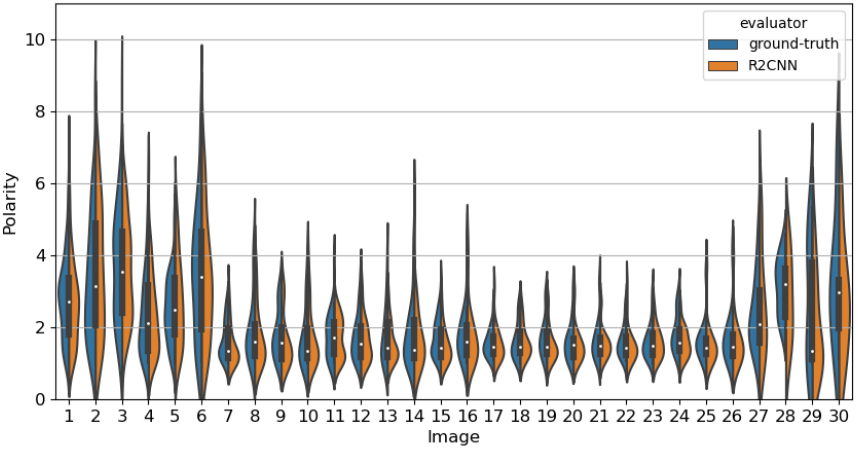
Cell polarities comparison on the *Test* data split.

#### 2) Cell polarity

Using OBBs instead of HBBs allows us to estimate the precise cell shape/polarity (i.e., no segmentation masks required). In fact, the cell shape for oriented and elongated cells simply cannot be estimated through HBBs, as illustrated in Fig. 1a. To determine whether the trained models could detect cell polarity differences comparably to human annotators, we performed a Kruskal-Wallis test on the distributions of polarities of each image on the *Annotators’ Comparison* dataset.

Fig. 7 illustrates cell polarity (i.e., OBB axis ratio) distributions in one of the images in the *Annotators’ Comparison* split of the dataset. A visual analysis indicates that all trained models performed similarly to human annotators in detecting changes in cell morphology. The best models were R3Det and R2CNN, which had no statistical difference with human annotators for all ten evaluated images from the *Annotators’ Comparison* dataset split. However, the worst-performing model (DCL) was still capable of scoring 7/10. Most models were indistinguishable from humans in most images as to their capacity to detect different cell polarities, except for one image. This image, shown in Fig. 8 (along with human annotations), illustrates several significant challenges in cell detection in bright-field images. More precisely, we can see some level of disagreement among human annotators both in OBBs orientation (Arrow 2), as well as which objects should be considered as cells (Arrow 1).

**Fig. 7:**
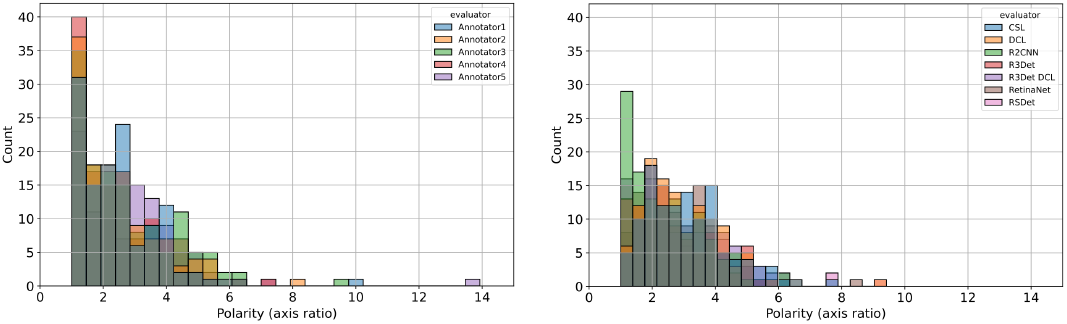
Example of polarity histograms (for image in Fig. 8). Bars represent the number of cells with a given polarity (left: human annotators; right: model results.

**Fig. 8:**
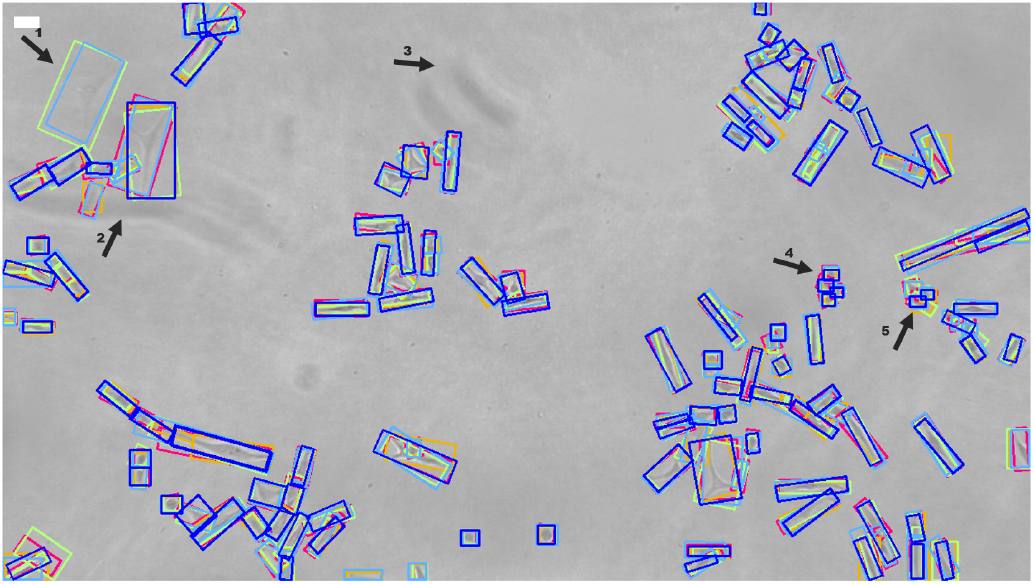
*Annotator’s Comparison* dataset image example. OBBs are colored according to each different human annotator. Arrows indicate challenging situations which might hinder the models’ performance, such as cells out of focus (1), blurry background (2, 3) and aggregated cells (4, 5). Scale bar (top-left white) equals 30 *μ*m.

We further explored the cell polarity distributions produced by all models in the *Test* data split, which contains a single annotation per OBB. Fig. 6 illustrates the distributions (kernel density estimations produced from polarities histograms) for ground-truth annotations and the results produced by R2CNN as an example model. Visually, model results are similar to human annotations overall. We then quantified the polarities error by calculating the Chi-distance for polarities distributions for all models, as shown in the last column of Tab. III. R3Det performed poorly compared to other models, presenting the highest Chi-distance value. Interestingly, this correlates with the confluence MRE values - in which R3Det also presented the highest error - since both estimations depend on the width and height of the OBBs. Meanwhile, similar to confluence estimation, the best models were RetinaNet and RSDet, respectively. As in the previous two biological applications, the best model for object detection (R2CNN) did not produce the smallest chi-squared distance for polarity estimation: *∼*12% higher than RetinaNet.

## VI. Conclusions

This work proposed the use of Oriented Bounding Boxes (OBBs) for the problem of cell detection in bright-field microscopy. We introduced an Oriented Cell Dataset (OCD) containing bright-field microscopy images with OBB and category (elongated or round) annotations. A subset of our data contains the labeling of five different experts, allowing us to perform Inter Annotator Analysis (IAA) and define suitable acceptance thresholds for evaluating OBB object detectors.

To corroborate our claim that OBBs (and OEs) are suitable representations for cell shapes, we presented a comparison of different annotation formats with the segmentation masks in the popular CTC dataset [12]. Our analysis showed that OBBs and OEs show considerable improvement over HBBs and AAEs, while being much cheaper to annotate than segmentation masks.

Our IAA analysis based on Krippendorff’s Alpha score showed a good/very good agreement among the human annotators despite the challenges involved with the dataset annotation process. In terms of cell shape annotation based on OBBs, our results indicated that the Intersection-over-Union (IoU) between two annotations of the same cell might not be close to the ideal value (one), which directly impacts the interpretation of classical object detection metrics. In particular, we show that using an IoU larger than 0.5 `is not feasible` for this dataset, whereas generic object detectors are evaluated with thresholds varying from 0.5 to 0.95 [29]. Using the suggested IoU threshold of 0.5, we concluded that R2CNN [17] presented the best mAP_50_ value among all tested OBB detectors.

Finally, we explored OBB detectors in the context of automating two important problems in cancer biology: cell confluence and polarity estimation. Regarding the two chosen applications, our results indicated that all models could estimate cell confluence in our dataset successfully, and most were statistically indistinguishable from humans in determining cell polarity. This enables the application of the proposed models as tools for automating cancer cell research. Additionally, our results indicate that the best model is not necessarily the same for distinct applications according to the corresponding task-specific evaluation metrics. Furthermore, they do not fully agree with the task-agnostic evaluation metrics commonly used for object detection. However, we emphasize that such findings were based on the *Test* split, which contains annotations from a single annotator, and could differ if compared to multiple human annotations. Despite this, all models achieved applicable results for the automation of cancer cell microscopy image analysis.

## Footnotes

1 Generated at Labsinal: Cell Signaling and Plasticity Laboratory – UFRGS (https://www.ufrgs.br/labsinal/)

2 Available at: https://ieee-dataport.org/documents/oriented-cell-dataset-ocd.

3 https://github.com/cgvict/roLabelImg.

4 http://celltrackingchallenge.net

5 https://opencv.org/

6 https://github.com/yangxue0827/RotationDetection

7 We used a Python package available at: https://pypi.org/project/kneed/

## References

[1] A. O. Silva, E. Dalsin, G. R. Onzi, E. C. Filippi-Chiela, and G. Lenz, “The regrowth kinetic of the surviving population is independent of acute and chronic responses to temozolomide in glioblastoma cell lines,” Experimental Cell Research, vol. 348, no. 2, pp. 177–183, 2016.

[2] S. Martinotti and E. Ranzato, “Scratch wound healing assay,” Epidermal cells: methods and protocols, pp. 225–229, 2020.

[3] J. T. Freitas, I. Jozic, and B. Bedogni, “Wound healing assay for melanoma cell migration,” Melanoma: Methods and Protocols, pp. 65–71, 2021.

[4] S. Busschots, S. O’Toole, J. J. O’Leary, and B. Stordal, “Non-invasive and non-destructive measurements of confluence in cultured adherent cell lines,” MethodsX, vol. 2, pp. 8–13, 2015.

[5] R. Paveley, N. Mansour, I. Hallyburton, A. Guidi, I. Gilbert, A. Hopkins, and Q. Bickle, “Whole organism high-content screening by label-free, image-based bayesian classification for parasitic diseases,” Plos Neglected Tropical Diseases, vol. 6, p. e1762, 2012.

[6] “Deep learning for computational cytology: A survey,” Medical Image Analysis, vol. 84, p. 102691, 2023.

[7] E. C. Filippi-Chiela, M. M. Oliveira, B. Jurkovski, S. M. Callegari-Jacques, V. D. d. Silva, and G. Lenz, “Nuclear morphometric analysis (nma): screening of senescence, apoptosis and nuclear irregularities,” 2012.

[8] A. M. Krebs, J. Mitschke, M. Lasierra Losada, O. Schmalhofer, M. Boerries, H. Busch, M. Boettcher, D. Mougiakakos, W. Reichardt, P. Bronsert, et al., “The emt-activator zeb1 is a key factor for cell plasticity and promotes metastasis in pancreatic cancer,” Nature cell biology, vol. 19, no. 5, pp. 518–529, 2017.

[9] Z. Zou, K. Chen, Z. Shi, Y. Guo, and J. Ye, “Object detection in 20 years: A survey,” Proceedings of the IEEE, 2023.

[10] J. Hayashida, K. Nishimura, and R. Bise, “Consistent cell tracking in multi-frames with spatio-temporal context by object-level warping loss,” in Proceedings of the IEEE/CVF Winter Conference on Applications of Computer Vision, pp. 1727–1736, 2022.

[11] L. N. Kirsten and C. R. Jung, “Cell tracking-by-detection using elliptical bounding boxes,” 2023.

[12] M. Maška, V. Ulman, D. Svoboda, P. Matula, P. Matula, C. Ederra, A. Urbiola, T. España, S. Venkatesan, D. M. Balak, et al., “A benchmark for comparison of cell tracking algorithms,” Bioinformatics, vol. 30, no. 11, pp. 1609–1617, 2014.

[13] M. Maška, V. Ulman, P. Delgado-Rodriguez, E. Gómez-de Mariscal, T. Nečasová, F. A. Guerrero Peña, T. I. Ren, E. M. Meyerowitz, T. Scherr, K. Löffler, et al., “The cell tracking challenge: 10 years of objective benchmarking,” Nature Methods, pp. 1–11, 2023.

[14] S. Ren, K. He, R. Girshick, and J. Sun, “Faster r-cnn: towards real-time object detection with region proposal networks,” IEEE transactions on pattern analysis and machine intelligence, vol. 39, no. 6, pp. 1137–1149, 2016.

[15] G.-S. Xia, X. Bai, J. Ding, Z. Zhu, S. Belongie, J. Luo, M. Datcu, M. Pelillo, and L. Zhang, “Dota: A large-scale dataset for object detection in aerial images,” in The IEEE Conference on Computer Vision and Pattern Recognition (CVPR), June 2018.

[16] X. Yang, G. Zhang, X. Yang, Y. Zhou, W. Wang, J. Tang, T. He, and J. Yan, “Detecting rotated objects as gaussian distributions and its 3-D generalization,” IEEE Transactions on Pattern Analysis and Machine Intelligence, 2022.

[17] Y. Jiang, X. Zhu, X. Wang, S. Yang, W. Li, H. Wang, P. Fu, and Z. Luo, “R2cnn: rotational region cnn for orientation robust scene text detection,” arXiv preprint 1706.09579, 2017.

[18] X. Yang, Q. Liu, J. Yan, A. Li, Z. Zhang, and G. Yu, “R3det: Refined single-stage detector with feature refinement for rotating object,” arXiv preprint 1908.05612, vol. 2, no. 4, 2019.

[19] W. Qian, X. Yang, S. Peng, Y. Guo, and J. Yan, “Learning modulated loss for rotated object detection,” arXiv preprint 1911.08299, 2019.

[20] T.-Y. Lin, P. Goyal, R. Girshick, K. He, and P. Dollár, “Focal loss for dense object detection,” in Proceedings of the IEEE international conference on computer vision, pp. 2980–2988, 2017.

[21] X. Yang, J. Yan, and T. He, “On the arbitrary-oriented object detection: Classification based approaches revisited,” arXiv preprint 2003.05597, 2020.

[22] X. Yang, L. Hou, Y. Zhou, W. Wang, and J. Yan, “Dense label encoding for boundary discontinuity free rotation detection,” in Proceedings of the IEEE/CVF Conference on Computer Vision and Pattern Recognition, pp. 15819–15829, 2021.

[23] D. Ker, S. Eom, S. Sanami, R. Bise, C. Pascale, Z. Yin, S.-i. Huh, E. Osuna-Highley, S. Junkers, C. Helfrich, et al., “Phase contrast timelapse microscopy datasets with automated and manual cell tracking annotations. sci. data 5, 180237,” 2018.

[24] S. Anjum and D. Gurari, “Ctmc: Cell tracking with mitosis detection dataset challenge,” in Proceedings of the IEEE/CVF Conference on Computer Vision and Pattern Recognition Workshops, pp. 982–983, 2020.

[25] J. Murrugarra-Llerena, L. N. Kirsten, and C. R. Jung, “Can we trust bounding box annotations for object detection?,” in Proceedings of the IEEE/CVF Conference on Computer Vision and Pattern Recognition, pp. 4813–4822, 2022.

[26] A. Angonezi, F. Oliveira, J. Faccioni, C. Cassel, D. Santos de Sousa, S. Vedovatto, G. Lenz, C. Jung, and L. Kirsten, “Oriented cell dataset (ocd),” 2024.

[27] T. T. D. Nguyen, H. Rezatofighi, B.-N. Vo, B.-T. Vo, S. Savarese, and I. Reid, “How trustworthy are performance evaluations for basic vision tasks?,” IEEE Transactions on Pattern Analysis and Machine Intelligence, 2022.

[28] J. Nassar, V. Pavon-Harr, M. Bosch, and I. McCulloh, “Assessing data quality of annotations with krippendorff alpha for applications in computer vision,” arXiv preprint 1912.10107, 2019.

[29] T.-Y. Lin, M. Maire, S. Belongie, J. Hays, P. Perona, D. Ramanan, P. Dollár, and C. L. Zitnick, “Microsoft coco: Common objects in context,” in Computer Vision–ECCV 2014: 13th European Conference, Zurich, Switzerland, September 6-12, 2014, Proceedings, Part V 13, pp. 740–755, Springer, 2014.

[30] K. He, X. Zhang, S. Ren, and J. Sun, “Deep residual learning for image recognition,” in Proceedings of the IEEE conference on computer vision and pattern recognition, pp. 770–778, 2016.

[31] J. Deng, W. Dong, R. Socher, L.-J. Li, K. Li, and L. Fei-Fei, “Imagenet: A large-scale hierarchical image database,” in 2009 IEEE conference on computer vision and pattern recognition, pp. 248–255, Ieee, 2009.

[32] A. F. Hayes and K. Krippendorff, “Answering the call for a standard reliability measure for coding data,” Communication methods and measures, vol. 1, no. 1, pp. 77–89, 2007.

[33] V. Satopaa, J. Albrecht, D. Irwin, and B. Raghavan, “Finding a “kneedle” in a haystack: Detecting knee points in system behavior,” in International conference on distributed computing systems workshops, pp. 166–171, IEEE, 2011.

[34] M. Everingham, L. Van Gool, C. K. Williams, J. Winn, and A. Zisserman, “The pascal visual object classes (voc) challenge,” International journal of computer vision, vol. 88, no. 2, pp. 303–338, 2010.

